# Functional Assessment of Cardiac Beat Dynamics Under Dynamic Flow: Insights from the Mera Microphysiological System

**DOI:** 10.64898/2026.05.20.726520

**Authors:** Nuno Almeida, Veasna Sum-Coffey, Patrick Costello, Conor Madden, Shane Devitt, Sumir Ramesh Mukkunda, Bhairavi Bengaluru Keshava, Sandra Sunil, Leon G. Riley, Seonaid Deely, Chiara Alessia de Benedictis, Mark Lyons, Finola E. Cliffe

## Abstract

Cardiac rhythm is a critical clinical indicator for cardiac arrhythmias and adverse events during drug toxicity studies. In vivo, cardiomyocyte responses to pharmacological agents occur within minutes and are strongly influenced by dynamic drug delivery through blood flow. However, conventional 2D and 3D static culture systems fail to replicate these fluid flow kinetics, limiting their physiological relevance for assessing beat rate responses.

Here, we present Mera, an advanced microphysiological system (MPS) developed by Hooke Bio, designed for high-throughput, long-term culture and functional analysis of 3D cardiac spheroids composed of human induced pluripotent stem cell-derived cardiomyocytes and cardiac fibroblasts. Mera enables dynamic perfusion, allowing investigation of cardiomyocyte beat rates under physiologically relevant flow conditions. The platform supports up to 640 spheroids per run and integrates automated imaging, fluid handling, and user-friendly software, operating under controlled physiological conditions (37°C, 5% CO₂). Flow rates are tunable between 0 and 12.5 mL/min to mimic in vivo environments.

Pharmacological testing with verapamil, isoproterenol, calcium chloride, and propranolol demonstrated real-time, reversible modulation of beat rate under flow, including recovery following drug-induced suppression. System variability was comparable to a temperature-controlled reference platform, supporting robust statistical analysis. Dose-response studies yielded IC₅₀ values consistent with literature, confirming physiological relevance.

Collectively, these results demonstrate that Mera provides a reproducible, scalable, and human-relevant platform for cardiac drug testing. By enabling dynamic drug exposure and automated analysis, Mera represents a powerful new approach methodology (NAM) for improving the predictive assessment of cardiac safety and beat-rate modulation drug responses.

## 1. Introduction

Cardiac rhythm and rate disturbances affects nearly 2% of the global population and is commonly treated with medications such as Verapamil (Vera) and Isoproterenol (IPT), with dosages tailored to each patient’s condition. Prior to market approval, all drug candidates undergo screening for potential cardiac side effects. However, despite advances in preclinical testing, a recent meta-analysis found that cardiac disorders remain the second leading cause of safety-related drug withdrawals (18.8%), following hepatotoxicity (27.1%). To reduce public health risks and costly late-stage failures, improving the predictive accuracy of cardiac liability assessments is essential to ensure that effective drugs advance through development (Craveiro et al., 2020).

Conventional approaches to studying cardiomyocytes in drug discovery often involve 2D cell cultures, animal models, or ex vivo tissue assays. Primary cardiomyocytes, commonly isolated from neonatal or adult rodents, are widely used to evaluate electrophysiological and contractile responses. However, these models suffer from limitations, including cellular immaturity, poor long-term viability, and lack of physiological complexity (Lundy et al., 2023). Additionally, human primary cultures are challenging to maintain and often lack the complex tissue architecture present *in vivo.* Rodent models, while widely used for cardiovascular studies, have notable interspecies differences, such as the absence of the hERG potassium channel in mice that limit their translational value (Vuorenpaa et al., 2023). Ethical considerations and poor predictability of human outcomes further constrain the utility of animal models.

More recently, stem cell-derived cardiomyocytes, such as those generated from induced pluripotent stem cells (iPSCs), have emerged as a more promising model for studying human-specific cardiac responses. Although iPSC-derived cardiomyocytes provide more human-relevant models for studying cardiomyocyte-related responses, there is currently no technology available that enables large-scale application for this approach.

Microphysiological systems (MPS) provide a solution to this challenge by closely capturing human physiology. Often named “organ-on-a-chip” technologies, MPS are advanced platforms that mimic the physiological environment of human organs or tissues *in vitro*. These systems combine human-derived cells, biomaterials, and microfluidic technologies to model the complex interactions and functions of organs at a micro-scale. MPS possess immense potential as an alternative to animal models in basic research, drug development, and personalised medicine. In cardiac research, MPS platforms are particularly valuable for preclinical drug development, as they provide a more accurate representation of human heart function and disease compared to traditional 2D cell cultures or animal models.

MPS can model a range of cardiac diseases, including ischemic heart disease, heart failure, and cardiomyopathies, offering a powerful tool for drug screening and disease modeling. These systems can also be integrated with other organ-on-a-chip models to study multi-organ interactions and better simulate the systemic effects of drugs (Ronaldson-Bouchard et al., 2022). The use of human-induced pluripotent stem cell (iPSC)-derived cardiomyocytes as 3D constructs in these systems allows for personalised models that can reflect individual genetic differences, enabling more precise testing of therapies, including those targeting genetic cardiac diseases or functional rhythm disturbances (Huh et al., 2010). 3D cardiac models can mimic critical physiological features such as the electrical activity of the heart, cellular interactions, tissue contractility, and the mechanical forces involved in heartbeats, making them essential tools for assessing drug effects on cardiac health (Liu et al., 2024). For example, worldwide projections are that cardiac rhythm abnormalities affect almost 2 percent of the global population (Kingma et al., 2023). Using MPS as platforms to host 3D tissues, drugs that alter beat rate or electrophysiological dynamics (QT interval changes) can be tested in a more physiologically relevant context, reducing the risk of adverse cardiac events in later clinical trials (Vuorenpaa et al., 2023; Liu et al., 2024).

In this study, we will showcase the effective use of the Mera system with beat modulation drug regimens that modulate the beat rate of cardiomyocytes. We tested two drug regimes: (1) Verapamil and calcium chloride (CaCl_2_), to model calcium channel blockade and rescue; and (2) Isoproterenol and propranolol (Propra), to demonstrate β-adrenergic modulation. A beat-rate modulating agent clinically used for controlling ventricular rate in supraventricular arrhythmias, Vera is a calcium channel blocker that promotes relaxation of vascular and arterial smooth muscle, resulting in vasodilation (Arefanian et al., 2023; Brozovich et al., 2016). It exerts its effects by inhibiting the L-type voltage-gated calcium channels, thereby blocking the influx of calcium ions across the cell membrane (Arefanian et al., 2023; Shah et al., 2022). While effective, calcium channel blockers can be highly toxic, and an overdose can lead to serious morbidity and even mortality (Hofer et al., 1993). Vera interferes with the rapid calcium influx into cardiac myocytes in the conduction system and smooth muscle cells of the vasculature, causing reduced myocardial contractility, prolonged conduction time, and vascular relaxation (Batalis et al., 2007). A common sign of verapamil toxicity is reduced beat rate (bradycardia), accompanied by hypotension, metabolic acidosis, and hyperglycemia (Farkhondeh and Mehrpour, 2020). In clinical practice, calcium chloride or calcium gluconate is often administered intravenously to treat calcium channel blocker toxicity, with both forms requiring intravenous (IV) delivery (Fahie and Cassagnol, 2024). This approach is based on the theory that an increased extracellular calcium concentration can promote calcium influx through the remaining functional L-type calcium channels (Chakraborty and Hamilton, 2024). This first regime was performed in the Mera system to model this effect, where cardiomyocyte beat rate is suppressed by Vera administration and subsequently restored following calcium supplementation.

The second regime employs IPT, which is a non-selective β-receptor agonist that activates both β1 and β2 receptors, leading to an increase in cardiac output, a reduction in peripheral vascular resistance, and relaxation of bronchial and gastrointestinal smooth muscles (Kieffer and Abel, 2016; O’Shaughnessy, 2012; Szymanski and Singh, 2024). As a result, isoproterenol is used in the treatment of heart block, cardiac arrest, asthma, and bronchospasm (Kieffer and Abel, 2016). β1 adrenergic receptors are postsynaptic receptors linked to G stimulatory proteins, primarily found in heart tissue (Bertam-Ralph and Amare, 2021). Activation of these receptors increases intracellular calcium levels, resulting in positive inotropy (enhanced contractility), positive lusitropy (improved relaxation), positive chronotropy (increased heart rate), and positive dromotropy (enhanced conduction velocity) (Szymanski and Singh, 2024). β2 adrenergic receptors are also postsynaptic and function similarly to β1 receptors. Their activation of G-protein-coupled receptors increases intracellular cyclic adenosine monophosphate, leading to bronchodilation and vasodilation in various arterial systems (Szymanski and Singh, 2024). Propra, a non-selective β-blocker, inhibits the action of catecholamines (adrenaline and noradrenaline) at both β1 and β2 adrenergic receptors (Srinivasan, 2019). By blocking these β receptors, propranolol prevents the sympathetic effects mediated through them (Srinivasan, 2019; Routledge and Shand, 1979). Although the IPT/Propra combination is not a standard clinical protocol, it effectively demonstrates controlled upregulation and downregulation of cardiomyocyte beat rate, highlighting the responsiveness of the Mera system to pharmacological modulation.

Additionally, we compared IC50 values for Verapamil across different cardiac spheroid models, showing consistency with published data from various 3D cardiac tissues. Notably, cardiomyocyte/cardiac fibroblast spheroids exhibited higher basal beat rates and IC50 values than cardiomyocyte-only spheroids, likely due to differences in measurement methods, Vera enantiomer potency, and cardiomyocyte maturation levels.

In this work, we report the use of Mera to modulate cardiomyocyte beat rate in a controlled and reproducible manner on a semi-automated, user-friendly MPS system. The objective of this work was to compare the Mera system in the maintenance of cardiomyocyte structure and function, as well as assess the response to various stimuli in the form of therapeutic cardiac drugs against regular *in vitro* cell culture studies. Overall, MPS’s allow a more human-relevant, ethical, and cost-effective alternative to traditional preclinical models, improving the efficiency and safety of drug development processes.

## 2. Materials and Methods

### 2.1. Chemical Sources

For cell maintenance and drug treatment, Dulbecco’s Modified Eagle Medium (DMEM) (no glucose no pyruvate, Gibco™ 11966025), endothelial cell growth media 2 (Promocell C-22011), dialyzed fetal bovine serum (Gibco™ A3382001), dulbecco’s phosphate buffered saline (D-PBS, Corning™ 15313581), gentamicin (Gibco™ 157500600), (±)-verapamil hydrochloride 99% (Thermo Scientific 329330010), and tween 20 blocking buffer, 1% in PBS (10X) (Thermo Scientific J61544AP) were obtained from Fisher Scientific (Dublin, Ireland). Galactose (Sigma G5388), sodium pyruvate (Sigma S8636), calcium chloride anhydrous, granular ≤7.0 mm ≥93.0% (Sigma C1016), isoproterenol hydrochloride (Sigma 420355), propranolol hydrochloride 99% (Sigma P0884), paraformaldehyde 95%, powder (Sigma 158127), bovine serum albumin heat shock fraction, pH 7, ≥98% (Sigma A9647), and glycine ≥99% (HPLC) (Sigma G7126) were purchased from Merck (Dublin, Ireland). Triton® X-100 (polyethyleneglycol tert-octylphenyl ether) molecular biology grade was purchased from VWR (Dublin, Ireland). Viability was assessed using the viability/cytotoxicity kit (Biotium 30002-T), purchased from VWR (Dublin, Ireland).

For immunofluorescence staining, primary antibodies were purchased from Invitrogen (California, USA) unless otherwise stated. Connexin 43 mouse monoclonal antibody (CX-1B1), Alexa Fluor™ 488 (138388, IF: 1:100), alpha-actinin-2 rabbit polyclonal antibody (Bioss Antibodies, bs-10367R-TR, IF: 1:200), cardiac troponin T, (MA1-16687: ICC: 1:150), mouse IgG1 kappa isotype control (14-4714-82, IF: 1:100), rabbit IgG isotype control (31235, IF: 1:200). Secondary Alexa 488 and 568 antibodies were purchased from Invitrogen (California, USA), and Alexa Fluor 568 Phalloidin (A12380) was acquired from Thermo Fisher Scientific (Waltham, USA). CytoVista™ 3D cell culture clearing reagent (Thermo Scientific V11326), readyProbes™ hydrophobic barrier pap pen (Thermo Scientific R3777), and proLong™ glass antifade mountant (Thermo Scientific P36982) were purchased from Fisher Scientific (Dublin, Ireland).

### 2.2. Cell culture

Cardiomyocyte spheroids at a density of 5000 cells/well were generated using 96-well V-bottom ULA plates (MS-9096VZ, PHC Europe B.V., Breda, The Netherlands) with iPSC-derived cardiomyocytes (iCell cardiomyocytes^2^ 01434, Fujifilm Cellular Dynamics, Inc. R1059) according to Fujifilm protocols. Spheroids were maintained either with cardiomyocyte maintenance media (Fujifilm Cellular Dynamics, Inc., M1003) or media containing DMEM, 10% dialyzed fetal bovine serum, 10 mM galactose, 1 mM sodium pyruvate, and 25 µg/mL gentamicin, and incubated in 5% CO₂ at 37°C. Half of the media was replaced every 2-3 days. Spheroids were ready to use in experiments after 7 days from plating, whereby regular beating was observed. Synchrony and onset of beating were assessed visually on a per-well basis. Cardiomyocyte and cardiac fibroblast spheroids (80%:20%) at a density of 5000 cells/well were generated using 96-well V-bottom ULA plates (MS-9096VZ, PHC Europe B.V., Breda, The Netherlands) with iPSC-derived cardiomyocytes and iPSC-derived cardiac fibroblasts (iCell cardiac fibroblasts 01434, Fujifilm Cellular Dynamics, Inc. cat. no. R1257). Cardiomyocyte and cardiac fibroblasts spheroids (80%:20%) were maintained either with cardiomyocyte maintenance media supplemented with co-culture supplements A and B (Fujifilm Cellular Dynamics) or media containing a mixture of 80% DMEM with 10% dialyzed fetal bovine serum, 10 mM galactose, 1 mM sodium pyruvate and 25 µg/mL gentamicin, with 20% endothelial cell growth media 2 (Promocell C-22011), and incubated in 5% CO_2_ at 37°C. Half of the media was replaced every 2-3 days. Spheroids were ready to use in experiments after 7 days from cell plating, whereby regular beating was observed. Synchrony and onset of beating were assessed visually on a per-well basis.

### 2.3. Immunofluorescence 2D cardiomyocytes staining

Immunofluorescence staining was performed as described in the protocols below. Cardiomyocytes were fixed in 4% paraformaldehyde (PFA) in phosphate-buffered saline (PBS) for 20 min at 4°C. Cells were permeabilized at room temperature (RT) with 0.1% Triton X-100 for 1 hour and then washed with PBS. To avoid nonspecific binding of antibodies, cells were incubated with blocking solution (BS): 3% BSA (Merck, Dublin, Ireland) in PBS. Next, cells were incubated overnight at 4°C with the primary antibodies. After overnight incubation, the unbound primary antibody was removed by performing three washing steps with PBS. Next, cells were incubated in the dark at RT for 1 hour with the secondary antibody diluted in BSA. In parallel, to label, identify, and quantify F-actin filaments within the cells, fluorescent phalloidin staining solution was added according to the manufacturer’s instructions. Cells were washed three times with PBS, and subsequently, cell nuclei were counterstained with 4,6-diamino-2-phenylindole (DAPI) (BD, New Jersey, US) at RT. A final washing step with 1X PBS was performed before acquiring the images at the Olympus IX70 fluorescence microscope.

### 2.4. Flow cytometry

Cardiac fibroblasts were stained for 20 min in the dark in the refrigerator in FACS buffer consisting of PBS with 2% FBS and 0.02% sodium azide. The following antibodies were used for staining: CD90 Antibody, anti-human, REAfinity™ (1:50) (Miltenyi Biotec, Germany), and APC Mouse IgG1 κ isotype control (1:50) (BD, New Jersey, US). Cells were analysed using a BD Accuri™ C6 Plus flow cytometer (BD Biosciences, Aalst, Belgium). Data were analysed using FACS DIVA software and with the free online software Floreada.io (https://floreada.io/).

### 2.5. Immunofluorescence 3D cardiomyocyte and cardiomyocyte/cardiac spheroid staining

Cardiomyocyte and cardiomyocyte/cardiac fibroblast spheroids were fixed in 4% PFA in PBS for 20 min at RT, followed by washing with PBS and incubation with 0.3 M glycine for 10 min at RT to quench the possible aldehydes from PFA fixation. After removing and rinsing the glycine with PBS, the spheroids were permeabilized overnight in 0.5% triton X-100. They were then incubated with a 3% BSA/0.1% tween 20/0.1 M glycine blocking solution in PBS for 2 hours at RT and subsequently overnight at 4°C with primary or conjugated antibodies in a 1% BSA/0.1% triton X-100 solution. Samples were then washed and incubated with secondary antibodies in a 1% BSA/0.1% triton X-100 solution overnight at 4°C. Cells were counterstained with DAPI for 2 hours at 4°C and cleared with cytoVista™ 3D cell culture clearing reagent (Thermo Scientific, Dublin, Ireland) for 2 hours. The cleared spheroids were then mounted on microscope glass slides, cat. no. 7107, inside a drawn square with a readyProbes™ hydrophobic barrier pap pen (Thermo Scientific, Dublin, Ireland) with proLong™ glass antifade mountant (Thermo Scientific, Dublin, Ireland). Images were captured using a Zeiss LSM 800 confocal. The following lenses were used for imaging: 10×/0.3 NA (dry), 20×/0.5 NA (dry). For excitation, diode lasers with wavelengths 405, 488, and 561 nm were used. Airy processing, maximal intensity orthoprojections, and export in TIFF format were performed using the original Zeiss ZEN software, with all parameters carefully adjusted.

### 2.6. Viability staining

Calcein AM (CAL-AM) and ethidium homodimer III (ETHDIII) were used to label viable/dead cells in cardiomyocyte spheroids. CAL-AM (2 µM) and ETHDIII (10 µM) were incubated for 3 hours before imaging, as presented in supplementary data, supplementary Fig. 5. Images were acquired using either the in-built custom microscope on Mera or the Olympus IX70 microscope in brightfield and fluorescence modes using FITC and Texas red filter sets.

### 2.7. Description and Device Fabrication/Assembly

Spheroids can be used to represent specific organs in the human body, while a microfluidic perfusion system such as Mera can be used to model fluid circulation in the human body. The concept of the MPS fluid flow is illustrated in Fig. 1, with (a) showing the position of the 3D spheroid and (b) the fluid flow recirculation through four interconnecting channels in the Bioplate. The focus of this paper is to demonstrate a scaled-back version of this concept by assessing a single and a co-culture cardiac model using a perfused MPS with a recirculation unit. The system comprises two main aspects: the Bioplate, which is essentially a culture plate, and the Mera system, which controls the perfusion through the Bioplate via recirculation and a series of pumps and valves.

**Fig. 1.**
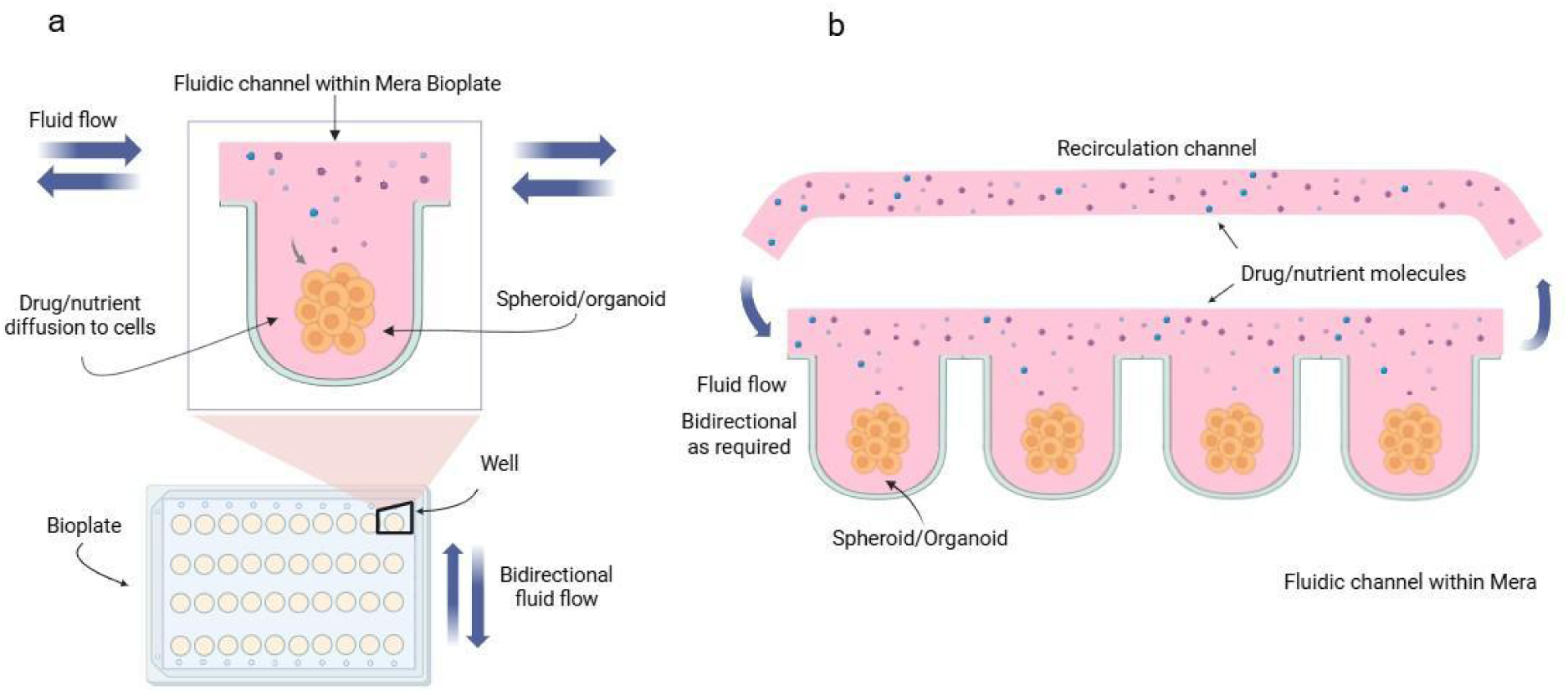
Illustration of Mera. a) Individual well in the Mera Bioplate showing the positioning of a 3D spheroid. b) Four interconnecting wells in the Bioplate. Created with BioRender.com

For all experiments using the Mera system, cardiomyocyte spheroids were transferred into Bioplate wells in 25 μL of media and then loaded onto the Mera fluidic assembly module as shown in Fig. 2. The fluidic assembly of Mera consists of four main parts: i) the lid, which holds the mounting for the recirculation assembly (Fig. 3). ii) the polytetrafluoroethylene (PTFE) membrane, iii) the Bioplate, which is arranged in a rectangular format consisting of a 10 x 4 grid containing round-bottomed wells into which spheroids can be situated, and iv) the fluidic module. To assemble the fluidic system, the PTFE plate is placed into the Bioplate, which is then inserted into the baseplate and then is mounted on top (Fig. 3). The PTFE plate provides the channel through which the fluid flows while providing suitable gaseous exchange and friction properties. The inlets and outlets of each given channel are found along the periphery of the Bioplate and form the entry and exit points for the fluid to run through the channels. Up to 40 spheroids can be cultured on a 4 x 10 well plate.

**Fig. 2.**
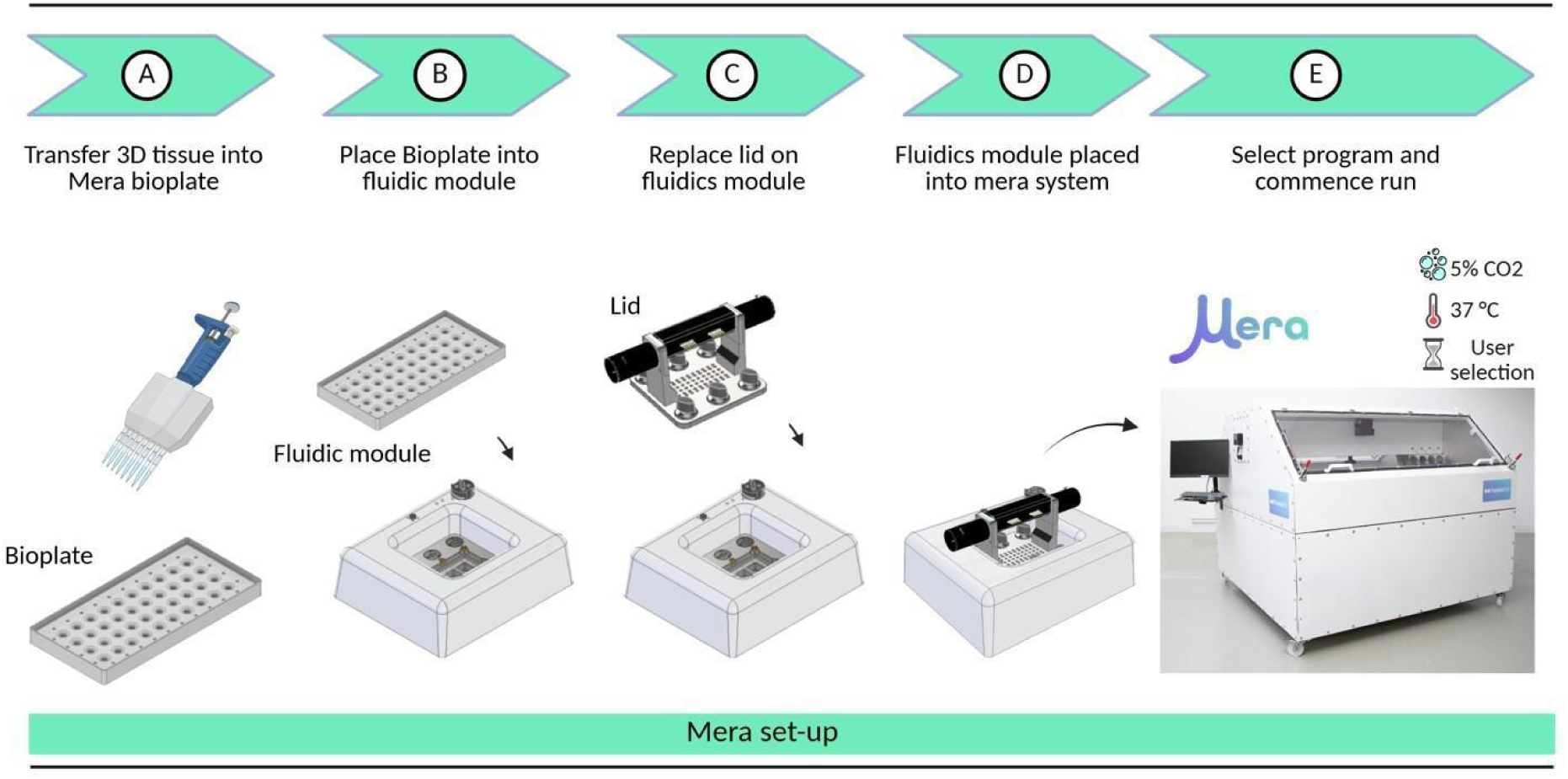
Mera setup including Bioplate and fluidic module loading. Steps A-D are undertaken in an aseptic environment such as a laminar flow hood. Created with BioRender.com

**Fig. 3.**
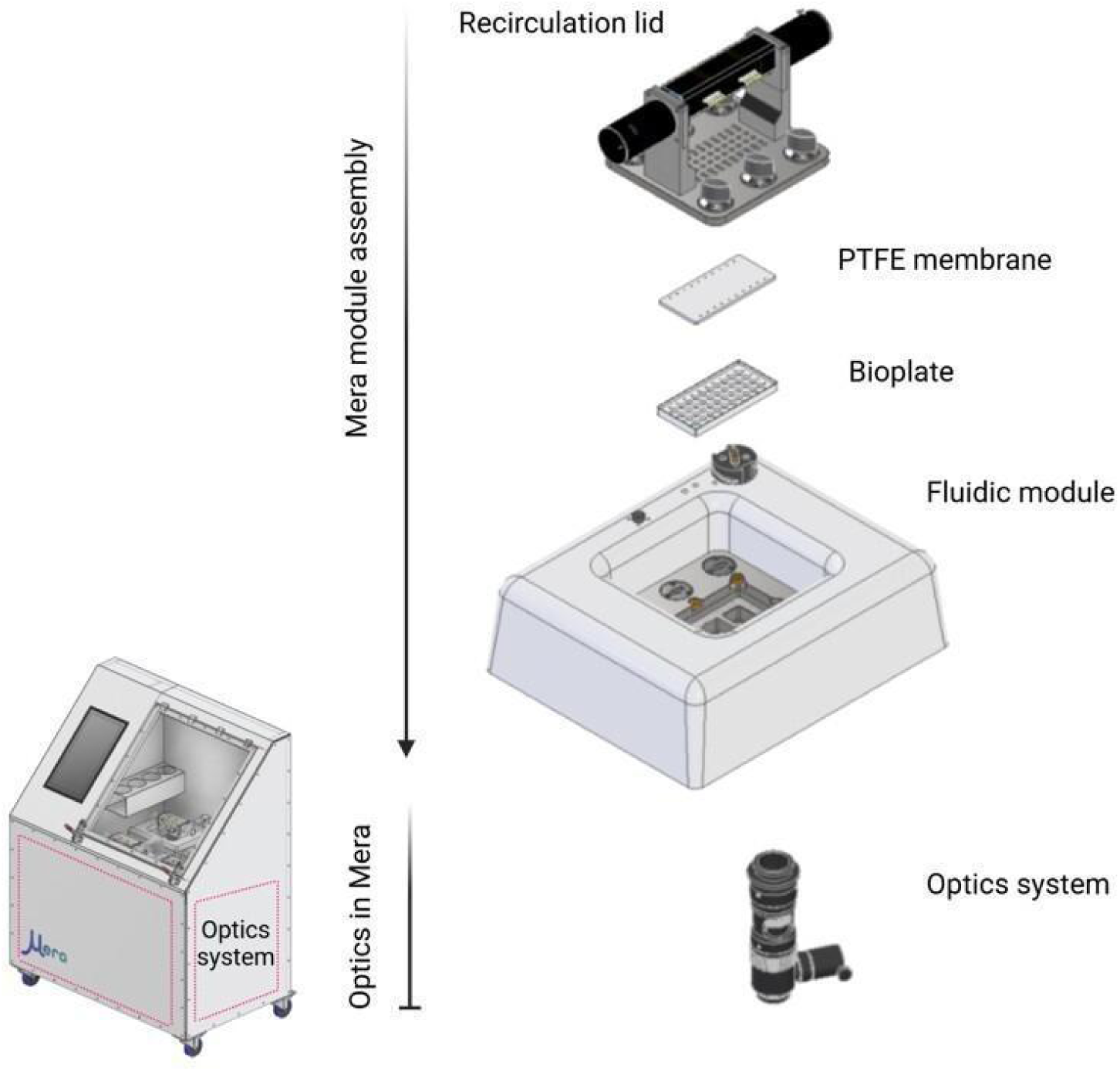
Exploded view of the Mera fluidic module with a stylized depiction of the location of the Mera optics system. The Bioplate is inserted into the baseplate within the fluidic module, the PTFE membrane is inserted to seal the channel, the lid is positioned onto the assembly and locked into place to provide a secure fluidic seal and sterile environment. After the module is placed into Mera, the integrated optics system can traverse between the four modules to capture images of the 3D tissues within the wells. Created with BioRender.com

### 2.8. Machining

All manufactured components were fabricated in-house as previously described [22].

### 2.9. Bioplate Design and Fabrication

The Bioplate consists of 40 (4 x 10 grid format) U-bottomed wells. The design of the Bioplate module has several advantages: 1) The dimensions and concave shape of the wells direct the spheroids to the center of the well to facilitate more accurate imaging and a consistent nutrient supply and decrease the possibility of damage due to shearing forces; 2) it also allows direct access for efficient loading of spheroids into the wells, manipulation during experiments, or removal for downstream analysis; 3) dead volume can be kept to a minimum; and 4) the optical transparency of the base of the wells enables high-quality optical readouts.

### 2.10. Bioplate integration with Mera

Mera consists of a central baseplate and associated lid (and PTFE plate) with optional recirculation functions into which the Bioplate sits. Perfusion into the Bioplate is supplied through the Fluidics Module via associated fluidic fittings and perfusion control valves. The Bioplate is integrated into the Fluidic module with the lid to create a complete assembly (Fig. 2), which houses the Bioplate whilst facilitating the control of the liquid supply through the mounted valves. It is designed with open channels to facilitate fluid flow while also providing a surface area to accommodate gaseous diffusion via osmosis, whilst constructing a channel within which fluid can pass. The Bioplate facilitates flow across four wells in a 10-channel format (4 wells x 10 channels, i.e., hosting spheroids of different organ types). Two pumps for recirculation are mounted on top of the lid, and the tubing connects the pumps to the lid. The fluid is directed around the baseplate and can be perfused 90 degrees upward through the internal geometry of the baseplate by opening the required valves, which facilitates the delivery of fluid to the Bioplate.

### 2.11. Perfusion control

A pump (Kamoer Fluid Tech, Shanghai, China) was used to control perfusion through the system. A constant rate of approximately 12.5 ml/min was used to prime and drug the spheroids hourly. This rate can be adapted depending on the individual needs of the cell model. A recirculation pump system (Takasago Electric Inc., Nagoya, Japan) was used for recirculating media.

### 2.12. Imaging system

A custom epi-fluorescence microscope was designed in-house and incorporated within the environmental chamber for automated in-situ imaging of spheroids. The microscope setup is previously described [22] with modifications to the light source. Briefly, the optical assembly of the imaging module consists of a light engine (SOLA, Lumencor, USA), a filter cube turret (X-FCR, Zaber, Canada), a camera tube lens (MTC00N, Zaber, Canada), filter cubes for brightfield, a 470 nm blue color (M470L5, Thorlabs, Germany), a 530 nm green color (M530L4, Thorlabs, Germany), and an objective lens with 5x magnification (46143, Mitutoyo, Japan). A complementary metal-oxide-semiconductor (CMOS) camera (acA1920-155ucMED, Basler AG, Germany) was used for microscopy imaging. The microscope was integrated with an automated 3-axis motion control system. A Python-based imaging GUI was developed using a basler pylon library to acquire the images and automatically control the motion of the microscope. This microscope was used to monitor spheroid placement within the wells of the Bioplate.

For reproducibility, brightfield and fluorescence images were acquired for all off-Mera samples using an Olympus IX70 inverted microscope (Olympus, Tokyo, Japan) with a 4x objective and an AmScope microscope digital camera (MU1003, AmScope, UK). Videos of cardiomyocyte beating were taken using the Mera optics in Mera maintained at 37°C. Video acquisition was taken before and after treatment with drugs, and the beat rate was manually counted.

### 2.13. Environmental Chamber

To grow and maintain reproducible spheroids, mammalian cells require a temperature of 37°C and 5% CO_2_ within a sterile environment. To achieve this, an environmental chamber with a dimension of 1.9 m x 1.9 m x 1.5 m was fabricated using stainless steel (SQ Fabs, Limerick, Ireland). The chamber was designed to be easily accessible for Bioplate removal, maintenance, and sterilisation procedures. A temperature control system, as previously described [22] maintained a temperature of 37°C. CO_2_ was supplied in a 6kg VB canister (BOC, Limerick, Ireland), which was safely tethered adjacent to the unit. CO_2_ sensor output is required to control the activation of the CO_2_ system as previously described.

### 2.14. Software and Electronics

The Mera module control system includes a set of printed circuit boards (PCBs). The PCBs are designed in-house with Altium PCB design software (Altium Limited, CA, USA). The PCBs are fabricated by JLCPCB (JiaLi Chuang Co., Limited, Hong Kong, China) and are assembled in-house.

### 2.15. Biological testing in Mera

For Bioplate loading, the fluidics module was detached from the Mera system and transferred to the laminar hood. Cardiomyocyte and cardiomyocyte/cardiac fibroblast spheroids were manually loaded into the Bioplate using an electronic pipette, and the Bioplate was then placed into its designated position within the fluidics module. The unit was sealed by securing the lid in place and tightening the lid connections. The exterior was disinfected with IPA spray and exposed to UV radiation for 15 min. Finally, the fluidics module was returned to the Mera system and reconnected to the electronic and fluidic lines. Several upstream processes (Fig 2., A-D) remain user-dependent, including manual placing of spheroids into the Bioplate, insertion of the Bioplate into the fluidic module, and subsequent placement of the module within the Mera system. The analytical workflow comprises automated data acquisition and live-cell imaging, while key analysis steps, including beat counting, were performed manually.

### 2.16. Temperature-controlled unit

A temperature-controlled system consisting of a plate holder, as displayed in Fig. 4, was manufactured in-house to facilitate imaging of the cardiomyocytes on the laboratory microscope (Olympus IX70). This system involves pumping heated water from the water bath (VWR® VWB2, VWR, Dublin, Ireland) through tubing to the internal fluid channels in the temperature control plate. The temperature-controlled plate was manufactured from aluminium, which allowed faster temperature convection to the microplate containing the cells. The pump controlled the fluid flow. Temperature control for the temperature incubation chamber was managed via a K-Type thermocouple (123-6306, Radionics, Dublin, Ireland) and an Analog Output K-Type thermocouple Amplifier – Breakout (AD8495) connected to an Arduino Uno (715-4081, Radionics, Dublin, Ireland). This measured the ambient air temperature above the wells. Code was written in the Arduino IDE to record the temperature of the thermocouple, to turn on and off the peristaltic pump, and to maintain the variable that acted as the set point for switching. A temperature set point measured by the thermocouple in the air of the chamber was used as a trigger to turn on or off the peristaltic pump. The set point was determined as follows: the system was calibrated by measuring liquid temperature in the wells of a 48-well plate and the surrounding air temperature in the chamber simultaneously. The temperature of the water bath and the pumping rate of the pump were adjusted (approximately 60°C) until the chamber could maintain a steady temperature of 37°C in the liquid of the wells of the 96-well plate. The equivalent air temperature in the chamber was used as a surrogate set point to turn on or off the pump when testing was performed with the plate containing cardiomyocytes.

**Fig. 4.**
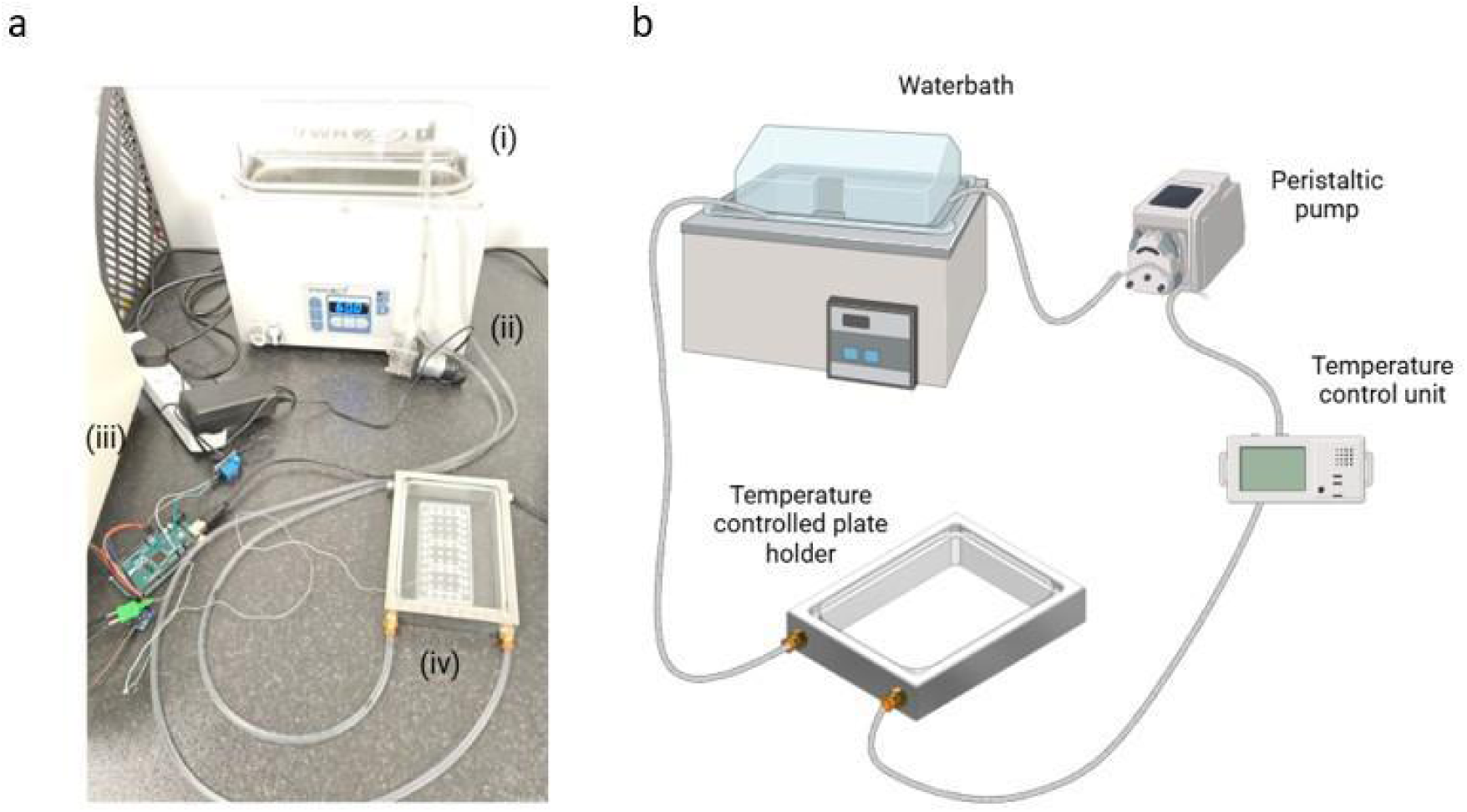
Temperature-controlled plate system. a) Picture of the layout of the system consisting of (i) waterbath, (ii) pump, (iii) fluid control unit and (iv) temperature-controlled plate holder. b) Schematic of the temperature-controlled plate system. Created with BioRender.com

### 2.17. Experimental Design

Cardiomyocyte spheroids were treated with drugs using different incubation conditions (as shown in Fig. 5) and then compared with basal control groups in the respective incubation conditions to assess drug response and viability of cardiomyocytes. The drug test groups were cardiomyocyte spheroids treated with either regime 1 (Fig. 5, top) or regime 2 (Fig. 5, bottom). For regime 1, spheroids were treated with 2 µM verapamil for 10 min and then washed with 2.5 mM calcium chloride and incubated for 1 hour. During regime 2, spheroids were treated with 100 µM isoproterenol hydrochloride for 10 min and then 100 µM propranolol hydrochloride for 10 min. Cardiomyocyte beat rates were recorded without any drug (basal beating), and in regime 1, 10 min after verapamil addition and 1 hour after calcium chloride (CaCl_2_) addition. For regime 2, cardiomyocyte beat rates were recorded without any drug (basal beating), 10 min after isoproterenol hydrochloride addition, and 10 min after propranolol addition. Each drug compound was delivered to the designated channel using an injection port. Each four-well channel was dosed sequentially to ensure precise measurement of cardiomyocyte beat rates.

**Fig. 5.**
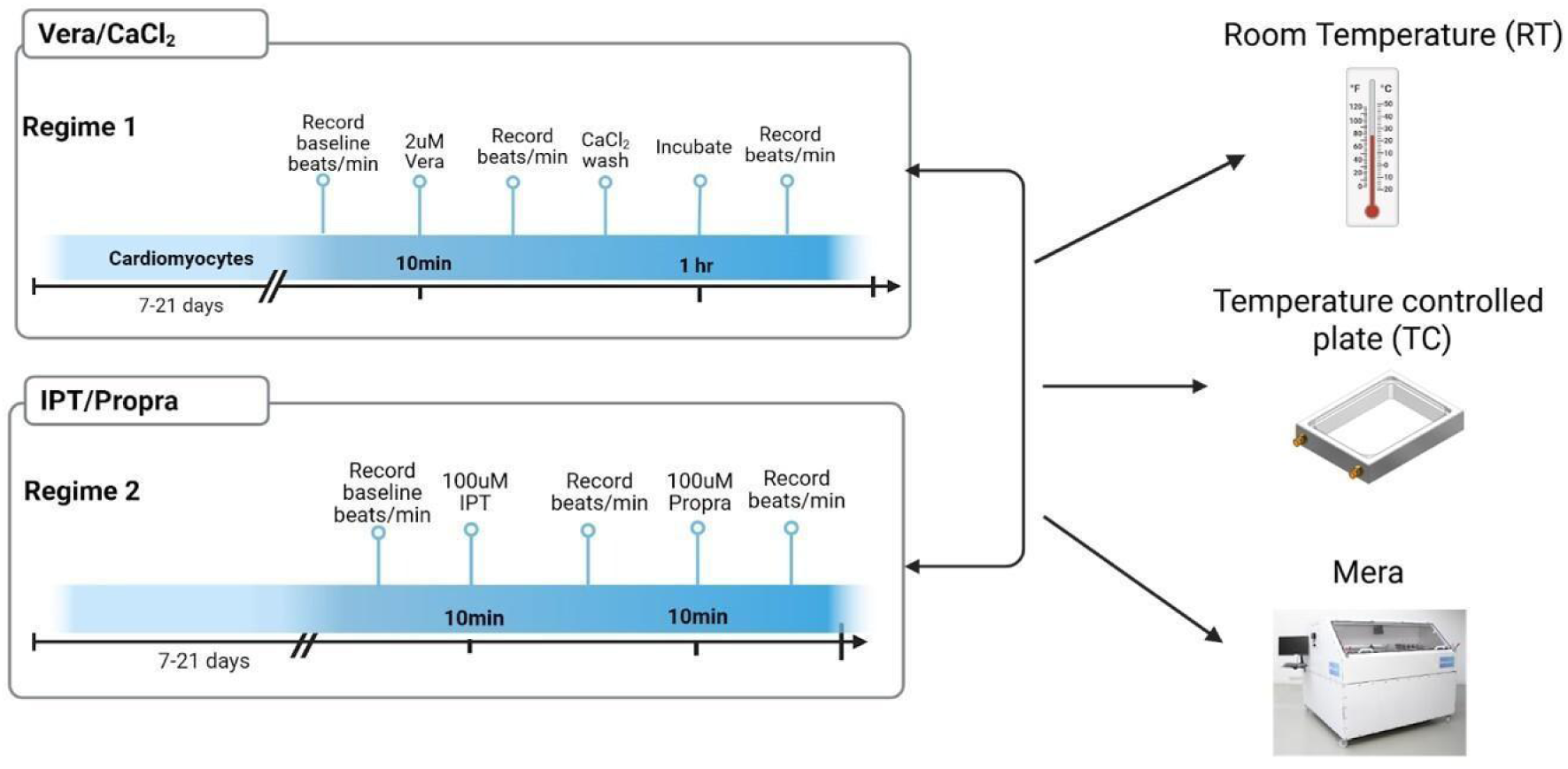
Experimental timeline. Cardiomyocytes were treated with Vera/CaCl_2_ and IPT/Propra, respectively, for each experimental condition, at room temperature, on the temperature-controlled plate, and in Mera. Created with BioRender.com

**Fig. 6.**
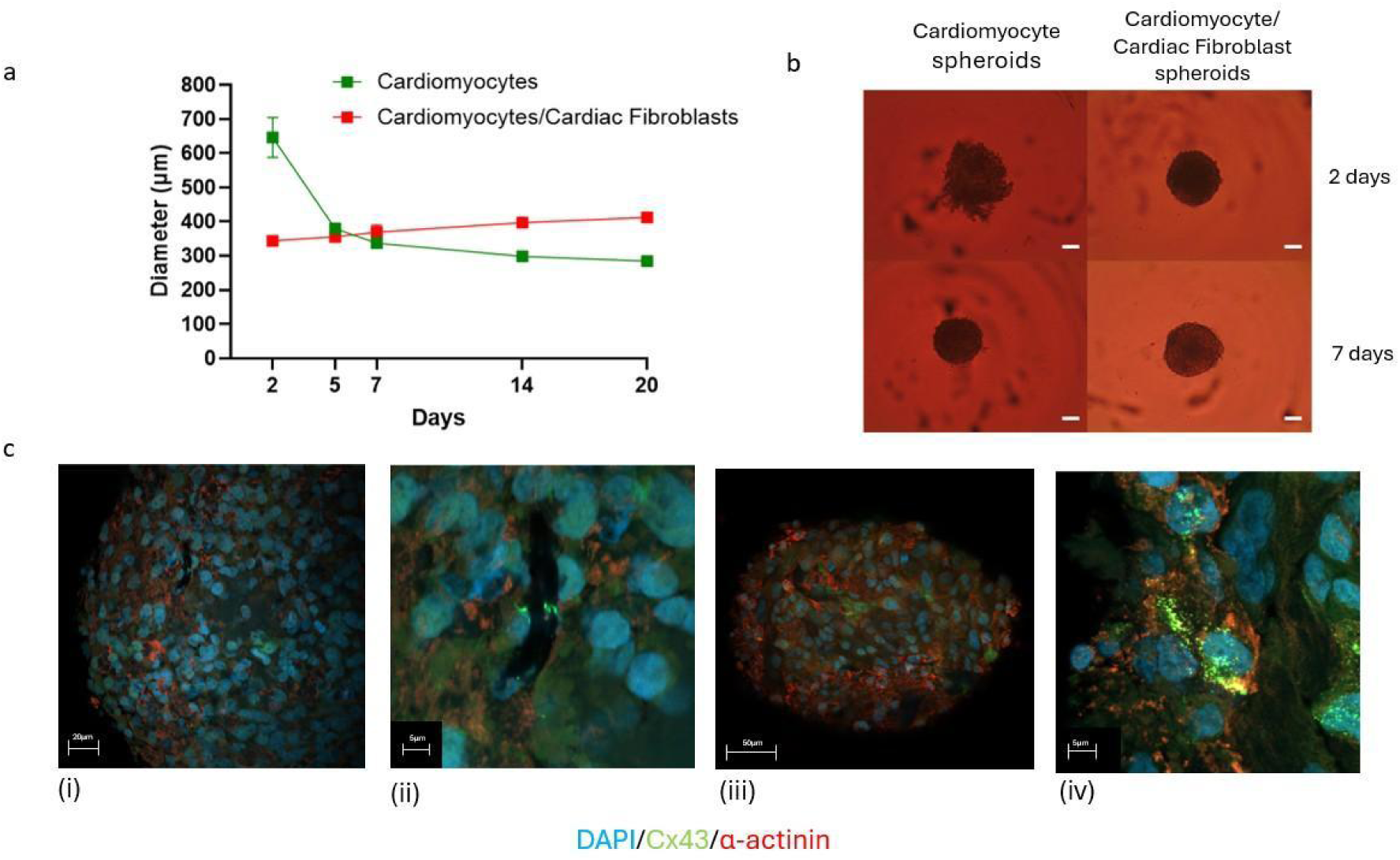
Characterisation of cardiomyocyte and cardiomyocyte/cardiac fibroblast spheroids. a) Growth profile of cardiomyocytes and cardiomyocyte/cardiac fibroblasts spheroids over 20 days. Cells were seeded at 5000 cells/well in a V-bottomed ULA plate, with three spheroids analysed per timepoint. b) Representative images of 3D cardiomyocytes and cardiomyocyte/cardiac fibroblasts spheroids after 48 h and 7 days of culture (scale bar = 100µm). c) Representative merged immunofluorescence staining of cardiomyocyte and cardiomyocyte/cardiac fibroblast spheroids with DAPI, Cx43, and α-actinin. i) & iii) overview picture of the spheroids captured with a 10×/0.3 NA (dry) lens ii) & iv) section of the spheroids captured with a 20×/0.5 NA (dry) lens

**Fig. 7.**
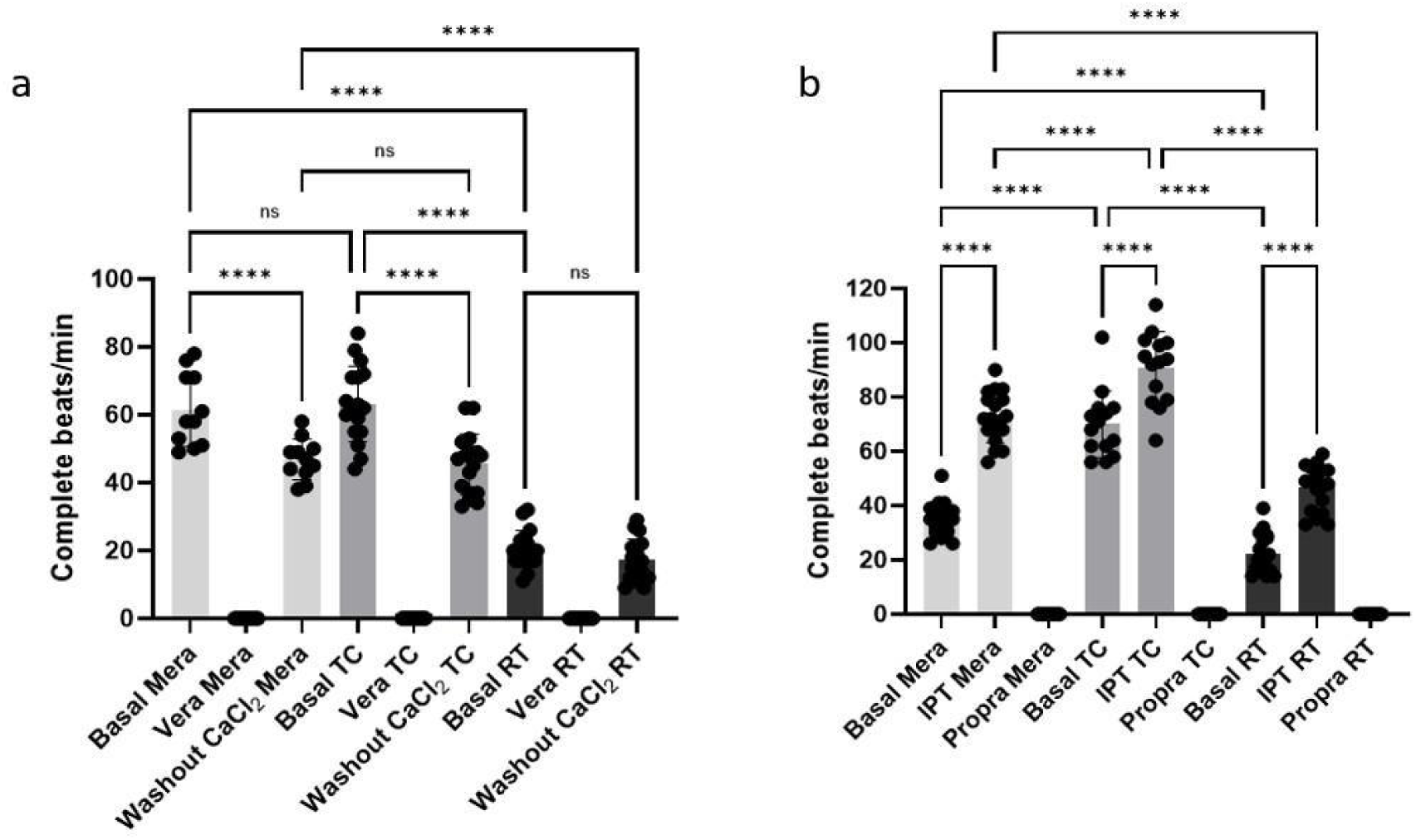
Impact of a) verapamil (Vera), calcium chloride (CaCl₂), and b) isoproterenol (IPT) and propranolol (Propra) exposure on the beating of cardiomyocyte spheroids on the Mera system, with an in-house temperature control device (TC) and at room temperature (RT). Beat rates were recorded before therapeutic incubation (basal), 10 min after Vera, IPT, or Propra addition, and 1 hour after CaCl₂. Results are expressed as the mean of complete beats/min ± SD, with the representation of each cardiomyocyte spheroid by a dot. At least three independent experiments were performed per condition, with a minimum of three spheroids analysed per experiment. For panel a), eleven spheroids were used in all Mera conditions, 15 spheroids in TC conditions and 16 spheroids in RT conditions. For panel b), 16 spheroids were analysed in all the Mera conditions, 11 spheroids in TC conditions and 18 spheroids in RT conditions. Significant p values are indicated by * in the individual Fig.s, where *p <0.05; **p <0.01; ***p <0.001 and ****p <0.0001.

The effects of drug treatment while using the Mera system were also compared to the incubation condition control groups, which included groups treated/not treated with the same drugs but (1) incubated in the temperature control unit (either the Basal/Drug TC group) or (2) incubated at room temperature (either the Basal/Drug RT group). For experiments using the Mera system, cardiomyocyte spheroids were transferred from 96-well V-bottom ULA plates (MS-9096VZ, PHCBI, Breda, The Netherlands) into Mera Bioplate wells containing DMEM and incubated at 37°C on the Mera system to determine the baseline cardiomyocyte, the beat rate in beats per minute (Basal Mera group), and the beats/minute in the test groups (Drug Mera). For room temperature experiments, cardiomyocyte beating was observed using an Olympus IX70 microscope. For experiments in the TC group, the TC unit was placed onto the microscope for imaging at 37°C. The liquid waste produced during the Mera experiments was collected in a designated glass waste receptacle and subsequently subjected to sterilisation via autoclaving at 121°C for 15 minutes under a pressure of 15 psi.

### 2.18. Drug Effect on Mera system compared to control groups, TC, and RT

For both drug dosing datasets, the data generated on Mera were compared to the data generated from the controlled plates (Basal RT group and Basal TC group). For one of the control groups, a second set of cardiomyocyte spheroids was placed in a standard 96-well round-bottom plate (MS-9096UZ, PHC Europe B.V., Breda, The Netherlands) and positioned within a custom temperature-controlling system as shown in Fig. 4, which could maintain a temperature of 37°C like the Mera system (TC group). The temperature control unit allowed for the maintenance of physiological temperature, like the Mera system, to evaluate the effect of temperature on the cardiomyocyte beat rate. The 96-well plates containing cardiomyocyte spheroids were placed in this temperature control unit after drug treatment or no drug treatment. Then, the cardiomyocyte beat rates were recorded, and comparisons between the control and test groups were assessed.

A third set of cardiomyocyte spheroids was treated/not treated with the drugs mentioned above and incubated in a standard 96-well round-bottom plate at room temperature (RT group) to assess the drug effect at room temperature, a second incubation control condition. For all test and control groups in both incubation conditions, identical dosing and washing procedures were applied to directly compare with the Mera groups.

### 2.19. Dose-response curves of Vera in vitro and on the Mera system

Dose-response curves for Vera were evaluated *in vitro* at RT as shown in Fig. 8. Vera was added to cardiomyocyte or cardiomyocyte/cardiac fibroblast spheroids from concentrations ranging from 0.01 µM to 5 µM as outlined in Fig. 8 (a), both *in vitro* at RT and in the Mera system, for 10 min. The beat rates were recorded for each concentration, plotted in GraphPad Prism (V10.5.0, GraphPad, MA, USA), and fitted in a sigmoidal 4PL curve to calculate the IC_50_ of each experimental group.

**Fig. 8.**
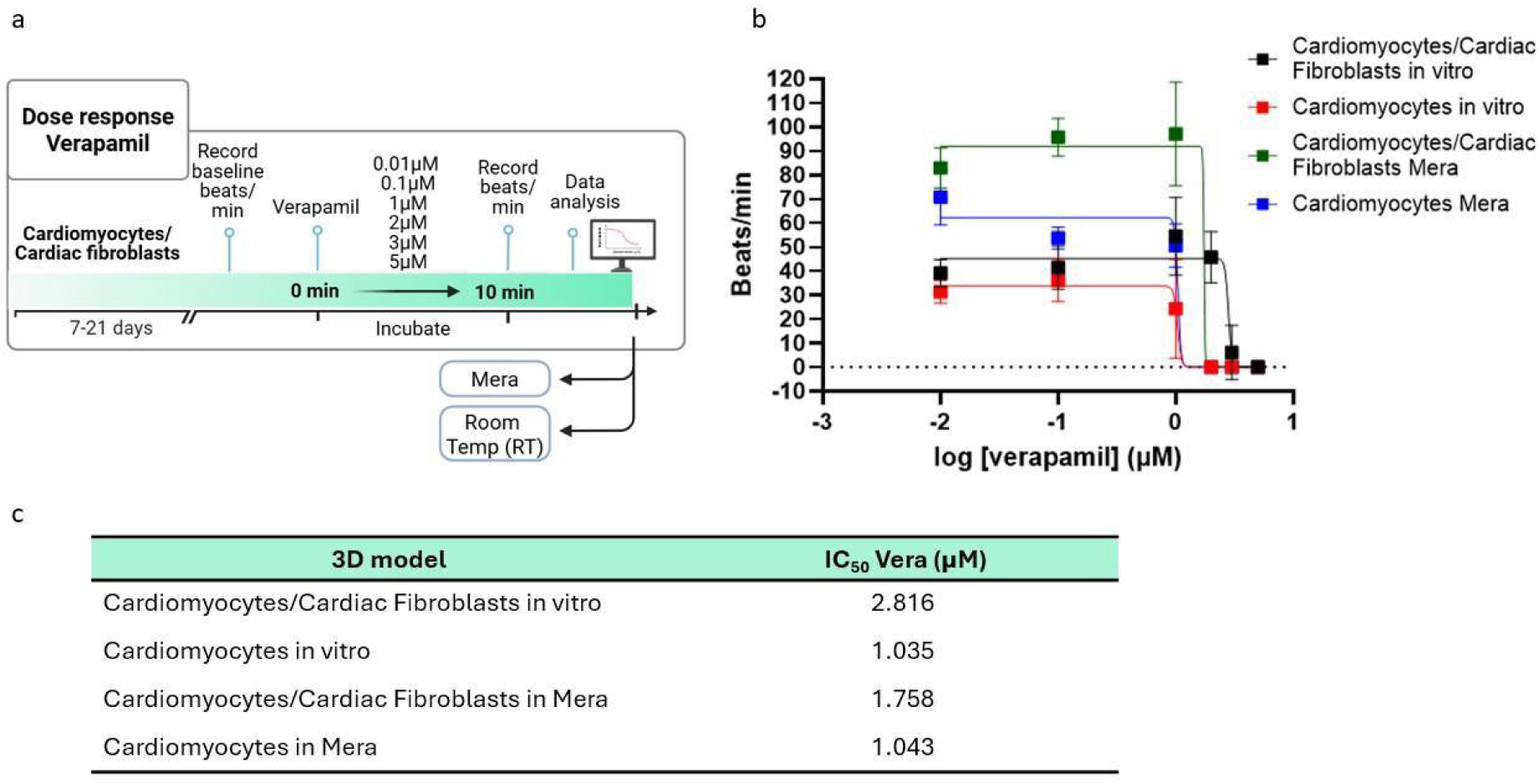
Dose-response analysis of Vera after 10 min incubation across a log-scale concentration range (0.01 - 5 μM). a) Experimental timeline created with BioRender.com, b) dose-response curves for cardiomyocyte and cardiomyocyte/cardiac fibroblast spheroids cultured *in vitro* and on the Mera system. c) Calculated Vera IC_50_ for cardiomyocyte and cardiomyocyte/cardiac fibroblast spheroids cultured *in vitro* and on the Mera system. Results are expressed as the mean of complete beats/min ± SD. At least three independent experiments were performed in each group, with eight spheroids analysed per condition.

### 2.20. Statistical Analysis

At least three independent experiments were performed for all the biological results unless stated otherwise. The data are presented as the mean ± standard deviation (SD). Statistical analyses were conducted using parametric tests (one-way ANOVA with Tukey’s multiple comparisons) as indicated in figure legends, following the determination of normal distribution by the Shapiro-Wilk normality test. p < 0.05 values were considered statistically significant. GraphPad Prism (GraphPad, MA, USA) was used for all the statistical analysis (V10.5.0).

## 3. Results

### 3.1. Characterisation of cardiomyocyte and cardiomyocyte/cardiac fibroblast spheroids

Before initiating biological testing, both cardiomyocytes and cardiomyocyte/cardiac spheroids were characterised, and spheroid viability was assessed on the Mera system over 48 h, as shown in Supplementary Fig. 5. Cardiomyocyte spheroids and cardiomyocyte/cardiac fibroblast spheroids were ready for use after five days of culture in 96-well V-bottomed ULA microplates. Fig. 6 (a) illustrates the growth profile of the spheroids over 20 days. Cardiomyocyte spheroids formed compact spheres by day 3-4 and, from day 5 onwards, maintained a diameter of approximately 300 µm throughout the 20- day culture period. Cardiomyocyte/cardiac fibroblast spheroids exhibited a different growth profile, reaching a diameter of approximately 400 µm by day 7, which was then maintained.Fig. 6 (b) highlights the faster compaction of cardiomyocyte/cardiac fibroblast spheroids compared to cardiomyocyte-only spheroids, as well as the larger size observed by day 7. The 80% cardiomyocytes:20% cardiac fibroblasts or 4:1 ratio, was chosen because it closely approximates the cellular composition of healthy cardiac tissue (Beauchamp et al., 2020).

A commonly used marker of cardiomyocytes is the sarcomere protein cardiac troponin T (Dewing et al., 2022). Immunohistochemistry staining was performed to detect the cardiac marker, troponin T, as shown in Supplementary Fig. 1. Cardiac fibroblasts were additionally assessed for CD90, a widely used marker for the identification of cardiac fibroblasts (Doppler et al., 2017), using flow cytometry (Supplementary Fig. 2). Cardiomyocyte and cardiomyocyte/cardiac fibroblast spheroids were stained for sarcomeric ⍺-actinin and connexin 43 (Cx43), as shown in Fig. 6 (c). Images of the separate channels and controls are provided in Supplementary Figures 3 and 4. Sarcomeric α-actinin, specifically the isoform α-actinin-2 (ACTN2), is an essential structural protein located at the Z-discs of sarcomeres in mature cardiomyocytes and plays a key role in maintaining scaromere integrity and function, the fundamental unit of muscle contraction (Hein et al., 2009; Fan et al., 2015; Lindholm et al., 2021). Immature cardiomyocytes exhibit a less organised and patchy sarcomere structure, while mature cardiomyocytes display a more organised and lengthier sarcomere network distributed across the entire cell (Skorska et al., 2022; Hsueh et al., 2023). Cx43 is a gap junction protein hemichannels in cardiomyocytes, enabling electrical signal transmission with synchronised contraction of the heart muscle and overall cardiac function (Boengler and Schulz, 2017). In addition, Cx43 plays a role in mediating cardiomyocyte maturation, establishing the interaction between cardiomyocytes and cardiac fibroblasts (Giacomelli et al., 2020). Our results show that cardiomyocyte/cardiac fibroblast spheroids exhibit a more organised sarcomere network compared to cardiomyocyte-only spheroids, as shown by the sarcomeric ⍺-actinin staining in Fig. 6 (c). Cx43 is also higher in cardiomyocyte/cardiac fibroblast spheroids compared to cardiomyocyte spheroids (Fig. 6 (c). Together, increased sarcomeric ⍺-actinin organisation and enhanced Cx43 expression indicate that cardiomyocyte/cardiac fibroblast spheroids represent a more mature cardiac model.

### 3.2. Impact of verapamil (Vera), calcium chloride (CaCl₂), isoproterenol (IPT) and propranolol (Propra) exposure on the beating of cardiomyocyte spheroids

A clear difference in beat rate was observed across all conditions when comparing RT with TC conditions or the Mera system, highlighting the importance of maintaining cardiomyocyte temperature as close as possible to physiological conditions (Figures 7a and b) (37°C). Previous studies using iPSC-derived cardiomyocytes have demonstrated that temperature influences beat rate, with increases in temperature directly proportional to a higher beat rate (Ikegami et al., 2023; Kanade et al., 2021; Rajan et al., 2020). The concentration of Vera used in this study has been previously reported in other iPSC cardiomyocyte studies (Sirenko et al., 2013; Seguret et al., 2024) to reduce the beat rate to 0 beats/min. As expected, (Fig. 7(a)), Vera decreased the cardiomyocyte beat rate in all conditions.

Recovery of cardiomyocyte beat rate was observed in the Mera system after Vera washout with CaCl₂, resembling the clinical outcome observed in patients who respond positively to calcium administration during verapamil overdose (Fig. 7(a)). CaCl₂ is used in nonacidotic patients since it delivers three times the amount of calcium as calcium gluconate, whereas calcium gluconate is preferred in acidotic patients, as CaCl₂ may exacerbate acidosis (Fahie and Cassagnol, 2024). Mera performed comparably to the temperature-controlled experiment, with both data sets demonstrating a significant difference between the basal cardiomyocyte beat rates and those observed after CaCl_2_ washout. β-adrenergic stimulation of cardiomyocyte spheroids with IPT increased the beat rate (Fig. 7(b)). In contrast, β-adrenergic antagonism of cardiomyocyte spheroids with propranolol arrested cardiomyocyte beating, even in the presence of isoproterenol (Fig. 7(b)). Additionally, as observed in regime 1, although basal beat rates between Mera and TC conditions differed significantly, similar trends were observed across all three experimental conditions. Notably, data reproducibility was significantly higher in the Mera system than in the TC unit (*p* = 0.018) with standard deviations of 8.05 and 11.42, respectively.

### 3.3. Vera IC_50_ for cardiomyocyte and cardiomyocyte/cardiac fibroblast spheroids *in vitro* and on the Mera system

We also demonstrate that the IC_50_ of Vera obtained using the Mera system is very similar to that measured under *in vitro* RT conditions, as shown in Fig. 8 (c). The cardiomyocyte/cardiac fibroblast spheroids exhibited a higher basal beat rate compared to cardiomyocyte-only spheroids, consistent with previous reports (Burnham et al., 2021).

## 4. Discussion

A similar profile of cardiomyocyte and cardiomyocyte/cardiac fibroblast spheroid formation has been observed before by others (Beauchamp et al., 2020; Arai et al., 2018; Burnham et al., 2021).The incorporation of cardiac fibroblasts with cardiomyocytes, among other cell types, showed the formation of improved sarcomeric structures, enhanced contractility, and more mature electrophysiological models than cardiomyocytes by themselves (Giacomelli et al., 2020). Additional strategies, including nanofibrous scaffolds with cardiomyocytes also showed an increased maturation status, with more sarcomeric ⍺-actinin staining (Iwon et al., 2024). Mechanical stretch can also play an important role in cardiomyocyte maturation through Cx43 signaling (Gu et al., 2021). Together, these findings support the importance of physiologically relevant multicellular and mechanically responsive environments for generating mature cardiac tissue models.

Vera, a phenylalkylamine class calcium channel blocker that has been used for more than half a century for the treatment of cardiovascular disease (Baky and Singh, 1982; Popovic et al., 2020). It is mainly prescribed for treating hypertension, arrhythmia, angina, and coronary artery disease (Arefanian et al., 2023; Koracevic et al., 2022) thus was selected as a clinically relevant beat-rate modulating compound for this study. Vera is widely distributed throughout the body tissues, and drug distribution to target organs and tissues is different with intravenous administration compared to oral administration, with about two-thirds of the drug being eliminated by hepatic metabolism, with excretion of inactive products in the urine and/or faeces (Hamann et al., 1984; Enna, 2007). After intravenous administration, the systemic clearance of Vera appears to approach liver blood flow. The high hepatic extraction results in low systemic bioavailability (20%) after oral drug administration (Hamann et al., 1984) while peak plasma concentrations are reached in 1 to 2 hours (Enna, 2007). An N-demethylated metabolite, norverapamil, which accumulates significantly in the blood, has been shown to have a fraction of the vasodilator effect of the parent compound in *in vitro* studies (Hamann et al., 1984; Enna, 2007). Vera can also block slow calcium channels in pancreatic beta cells and inhibit insulin release, thereby causing hyperglycemia (Louters et al., 2010). Bradycardia and hypotension are the most serious complications from a verapamil overdose, as both can lead to death if the patient is left untreated (Fahie and Cassagnol, 2024).

The binding of Vera was found to be specific, saturable, and reversible (Garcia et al., 1984), supporting the rationale for drug washout and subsequent calcium administration in the present study. Since no specific antidote exists for verapamil toxicity, removing the drug from the gastrointestinal tract is essential, and intravenous calcium should be administered to symptomatic patients. (Hofer et al., 1993; St-Onge et al., 2017). The rationale for calcium administration is that elevating extracellular calcium concentrations facilitates calcium influx through unblocked L-type calcium channels (Chakraborty and Hamilton, 2024).

In this study, the fluidic recirculation and injection capabilities of the Mera platform enabled modelling of both verapamil exposure and calcium rescue under dynamically perfused conditions. This represents an advantage over conventional static systems by enabling closer simulation of drug removal and therapeutic intervention workflows observed in vivo. Treatment of verapamil overdose with intravenous calcium (CaCl₂ or calcium gluconate) administration has been reported successfully in some patients (Barrow et al., 1994; Lipman et al., 1982) but also without response or negative outcomes in others (Atemnkeng et al., 2021; Crump et al., 1982; Sim and Stevenson, 2008).

IPT (also known as isoprenaline) β-adrenergic agonist clinically used for the treatment of bradycardia and, in some cases, complete heart block, β-blocker overdose, and cardiogenic shock-associated hypotension (O’Shaughnessy, 2012). Oral IPT is well absorbed but is subjected to first-pass metabolism, with only 4% of the drug being available (George, 1981; Redwood, 1969). Intravenous administration of IPT is 1,000 times more potent than oral administration (Conolly et al., 1972). Its plasma half-life is around 2.5 to 5 hours, and it is mainly metabolized by conjugation in the liver and lungs and is excreted in urine (Motwani and Saunders, 2024). The results observed with IPT on Mera are in line with what has been shown in the literature previously in studies using iPSC cardiomyocytes (Sirenko et al., 2013; Goldfracht et al., 2019; Grimm et al., 2018).

Propra is a non-specific β1/β2-adrenergic receptor antagonist that is mainly prescribed for cardiac disorders but also shows multiple beneficial off-target therapeutic effects in other pathologies (Cuesta et al., 2022). Propra is highly lipophilic, and administration can be either oral or intravenous (Shahrokhi and Gupta, 2024). Following oral administration, the drug is completely absorbed, with most of the dosage being removed via hepatic extraction, with only 25% of the drug reaching systemic circulation due to the first-pass metabolism in the hepatic circulation (Shahrokhi and Gupta, 2024; Shin and Johnson, 2007). It is extensively metabolized, and most of its metabolites are excreted in urine (Shahrokhi and Gupta, 2024; Shin and Johnson, 2007). The plasma half-life of Propra is between 3 to 6 hours (Srinivasan, 2019; Routledge and Shand, 1979). The active metabolite of propranolol is 4-hydroxypropranolol, which is formed through hydroxylation using the CYP2D6 enzyme (Shahrokhi and Gupta, 2024; Ripley and Saseen, 2014).

Other studies have already shown a decrease in beat rate in iPSC cardiomyocytes treated with Propra (Sirenko et al., 2013; Grimm et al., 2018). Notably, Grimm et al. utilized cardiomyocytes derived from the same donor source as those used in this study and reported comparable responses to both isoproterenol and propranolol exposure (Grimm et al., 2018), further supporting the reproducibility and translational relevance of the present findings.

The IC50 values obtained for Vera treatment in this study were comparable to values previously reported in engineered human cardiac tissue models. Turnbull et al. obtained an IC_50_=0.61 µM for Vera using a model of human embryonic stem cell-derived cardiomyocytes mixed with collagen and cultured on force-sensing elastomer devices (Turnbull et al., 2014). Mannhardt et al. reported an IC50 of around 0.3 µM for Vera treatment using a fibrin-based human-engineered tissue containing cardiomyocytes in agarose casting molds with solid silicone (Mannhardt et al., 2016). Seguret et al. showed an IC_50_=0.677 µM upon Vera treatment in a 3D ring-shaped cardiac tissue model generated from human induced pluripotent stem cell-derived cardiomyocytes and human dermal fibroblasts (Seguret et al., 2024). Our IC_50_ values for Vera treatment in cardiomyocyte spheroids are close to those values in both in vitro and Mera experiments, as observed in Fig. 8 (c), with the cardiomyocytes/cardiac fibroblast spheroids having higher values both in vitro and in Mera.

Multiple factors can explain the differences in the values obtained. First, the methods used to calculate the IC_50_ values of Vera are different between the authors, as we used the effect of Vera on complete beats/min to calculate the IC_50,_ and others used different parameters, like the contraction amplitude (Seguret et al., 2024; Turnbull et al., 2014; Mannhardt et al., 2016). Second, the different Vera enantiomers can also affect the IC_50_ values, with (-)-Vera being more potent than (+)-Vera in terms of its negative inotropic effect (Ferry et al., 1985). Finally, the maturation level of the cardiomyocyte can affect the response to drugs, as long-term cultured cardiomyocytes, 3D engineered cardiac tissues, and electrically stimulated cardiomyocytes demonstrated less sensitivity to Vera (Navarrete et al., 2013; Mathur et al., 2015; Li et al., 2024). This can explain the higher IC_50_ values obtained for our co-culture model.

Overall, this study highlights the variability in the basal beat rate of cardiomyocytes across experimental batches, providing a plausible explanation for the variations observed in basal beat rates between experiments. It also demonstrates the potential of Mera to effectively simulate therapies used in clinical settings and obtain accurate dose-responses.

Liu et al., provided a comprehensive review of heart-on-a-chip platforms, underscoring their capacity to recapitulate key aspects of human cardiac physiology for applications in disease modelling and pharmacological testing (Liu et al., 2025). Complementing this, Wang et al., discuss the development of microfluidic systems tailored for real-time monitoring of cardiomyocyte electromechanical activity, highlighting the integration of sensing technologies to assess functional parameters such as contractility and electrophysiology (Wang et al., 2025). Oleaga et al., have successfully predicted human cardiac toxicity upon compound transformation through hepatic metabolism in a multi-organ microfluidic device (Oleaga et al., 2018). Additional work by Cofiño-Fabres et al., demonstrated the fabrication of 3D heterotypic human cardiac tissue-on-chip constructs under flow conditions, revealing enhanced contractile performance, improved electrophysiological maturation, and endothelial–cardiomyocyte crosstalk (Cofiño-Fabres et al., 2024). While more recently, Liu et al. developed a tri-culture heart-on-a-chip model incorporating iPSC-derived cardiomyocytes, fibroblasts, and vascular endothelial cells under dynamic perfusion. This system enabled the formation of aligned multilayered 3D cardiac tissue that remained viable and functionally stable for over 60 days (Liu et al., 2024). Notably, endothelial cells exposed to flow exhibited increased barrier integrity, while tri-culture conditions upregulated the ventricular-specific marker IRX4 and significantly enhanced contractility compared to bi-culture models. Collectively, these studies highlight the increasing biological complexity and physiological relevance of cardiac MPS platforms, underscoring their growing importance in cardiovascular research and therapeutic development.

While our MPS model proved useful for the assessment of cardiomyocyte and cardiomyocyte/cardiac fibroblast spheroid responses to beat modulator compounds, particularly in terms of speed and cost-effectiveness, there are some limitations of the study that should be acknowledged. The current model is very simple and would benefit from increased complexity by incorporating additional cardiac cell types, such as endothelial and immune cells. It would also be advantageous to evaluate to a full extent the effects of cardiac drugs that act on several targets on other organ systems (e.g., liver, kidneys, etc.). Beat rate assessment provides a simple, high-throughput method for evaluating cardiomyocyte activity, but it has significant limitations when compared to more comprehensive functional readouts like electrophysiology, contractility, and calcium transient analyses (Blazeski et al., 2012; Sirenko et al., 2013; Dou et al., 2022; Vuorenpaa et al., 2023). For example, beat rate measurements alone do not provide information on contractile force or relaxation kinetics, both o f which are important parameters for assessing heart failure or drug-induced cardiotoxicity (Sirenko et al., 2013; Laurila et al., 2016; Hinata et al., 2022). Changes in beat rate can mask or distort underlying electrophysiological issues, making it difficult to determine if a drug effect is direct or merely a consequence of the altered rate (Laurila et al., 2016; Mannhardt et al., 2017). Beat rate assessment by itself does not allow the distinction between different cellular mechanisms in cardiac pathologies; for example, it cannot pinpoint whether an arrhythmia is caused by specific ion channel blocks or by defects in calcium handling, whereas electrophysiology and calcium imaging can (Laurila et al., 2016;Wells et al., 2019; Burnham et al., 2021). Additionally, video-based beat rate detection can be sensitive to movement artifacts and may fail to accurately record very fast beating or brief pauses (arrhythmic events) that more precise methods like patch clamp or multi-electrode arrays (MEAs) would capture (Laurila et al., 2016; Ito et al., 2020; Dou et al. 2022).

Currently cardiac models are not the primary focus of most commercial organ-on-a-chip or MPS developers, with more focus placed on hepatocyte models. However, advances are being made with multi-organ systems that combine hepatocytes and cardiovascular models to assess toxicity. Oleaga et al. have successfully predicted human cardiac toxicity upon compound transformation through hepatic metabolism in a multi-organ microfluidic device (Oleaga et al., 2018). Such approaches highlight the future potential of interconnected MPS platforms for more comprehensive evaluation of drug efficacy, metabolism, and systemic toxicity.

Cardiac MPS have advanced considerably in their ability to generate high-resolution functional outputs, particularly electrophysiological activity and contractility. Platforms incorporating microelectrode arrays (MEAs) are now well established for quantifying field potentials, conduction velocity, and arrhythmogenic events, and are widely used in preclinical cardiotoxicity assessment, including studies aligned with the Comprehensive in vitro Proarrhythmia Assay (CiPA) framework.

More recently, microfluidic heart-on-chip systems have integrated electrical and mechanical sensing with controlled perfusion and tissue architecture, enabling simultaneous assessment of electrophysiology, contractility, and microenvironmental cues. Commercial platforms such as those developed by BiomimX (e.g., uHeart) exemplify this approach, providing 3D cardiac tissues with embedded electrophysiological readouts, microfluidic control, and mechanical stimulation to recapitulate physiologically relevant cardiac function. This configuration enables measurement of field potentials, beat rate, and arrhythmic events under pharmacological perturbation. Notably, this system has demonstrated predictive capacity for QT interval prolongation and pro-arrhythmic risk, achieving high sensitivity and specificity relative to clinical outcomes, thereby underscoring its translational utility in preclinical safety pharmacology (Visone et. al. 2023; Visone et al. 2024)

In parallel, multi-organ MPS platforms from Hesperos extend this capability by incorporating cardiac modules within interconnected systems, enabling evaluation of systemic and metabolism-mediated cardiotoxicity over longer culture durations. Cardiac function is assessed using non-invasive, engineering-based readouts, including measurements of contractile force, beat frequency, and field potential duration (as a surrogate for the QT interval).

MIMETAS’s OrganoPlate system supports perfused three-dimensional (3D) endothelial microtissues under controlled flow, enabling the recreation of physiologically relevant haemodynamic conditions such as shear stress. These models have been widely applied to investigate angiogenesis, endothelial barrier function, and vascular dysfunction, all of which are central to cardiovascular disease pathophysiology. Although not intrinsically designed for electrophysiological or contractility measurements, the OrganoPlate is frequently combined with external analytical tools to extend functional cardiac assessment (Chapman et. al. 2024).

Incorporating recent studies, Emulate’s organ-on-chip platforms provide a mechanistically informative NAM approach for cardiovascular toxicity and disease modelling. Multi-lineage heart-chip models integrating iPSC-derived cardiomyocytes and endothelial cells under dynamic flow and cyclic strain have been shown to enhance cellular maturation and functional stability while enabling assessment of drug-induced cardiotoxicity across both myocardial and vascular compartments (Hou et. al. 2026). These systems demonstrate predictive capability for complex toxicities, including cancer therapy–associated cardiovascular effects. Complementary vascular organ-chip models, including a human coronary artery tri-culture system, further recapitulate key features of vascular inflammation and mechanobiology, such as the anti-inflammatory effects of pulsatile mechanical strain and multicellular interactions within the arterial wall. Collectively, these studies highlight the capacity of Emulate’s microphysiological systems to integrate haemodynamic forces, multicellular architecture, and functional readouts, enabling translational assessment of cardiotoxicity and vascular disease processes within a unified, human-relevant platform (Hou et. al.).

The Mera platform is best positioned as a scalable, modular MPS with strengths in throughput, flexibility, and system integration, rather than as a fully integrated high-content cardiac readout platform. In contrast to established cardiac MPS technologies, such as those developed by BiomimX and Emulate, which incorporate embedded electrophysiological and mechanical sensing for high-resolution functional characterisation, Mera prioritises experimental scalability and parallelisation. This architectural distinction enables higher-throughput experimental designs and facilitates combinatorial testing across multiple conditions or tissue types, aligning more closely with early-stage screening and systems-level interrogation. While multi-organ platforms such as those from Hesperos similarly support integrated biology, they typically operate at lower throughput and with greater experimental complexity However, Mera currently lacks integrated, real-time functional readouts such as microelectrode array-based electrophysiology or direct contractility measurements, which are key features of more mature cardiac MPS platforms. As a result, its application to detailed cardiac functional assessment remains dependent on external analytical methods. Future integration of embedded sensing modalities represents a logical next step to enhance its utility in cardiotoxicity screening and mechanistic cardiac studies.

Despite these limitations, Mera offers several distinct design advantages relative to existing MPS platforms. Notably, the system is engineered without the use of polydimethylsiloxane (PDMS), a material widely employed in organ-on-chip devices but known to exhibit significant absorption of small hydrophobic molecules, potentially confounding drug exposure and pharmacokinetic interpretation. In contrast, materials used in Mera were selected based on stability, biocompatibility, and compatibility with standard sterilisation protocols, supporting reproducibility and translational relevance. In addition, the platform is designed to operate with reduced fluid volumes, improving cost efficiency while more closely approximating physiological ratios of drug and analyte concentrations. This is particularly relevant for studies where accurate exposure dynamics are critical.

The platform has been manufactured and tested, and in its current configuration can support the concurrent culture of up to 640 three-dimensional tissues or spheroids. This capacity is enabled by a modular architecture comprising 16 fluidic modules, each housing a 40-well Bioplate. This design allows either full-system operation or selective use of individual modules, providing experimental flexibility. Such throughput facilitates the evaluation of broader dose ranges and increased technical replicates, which are important considerations for early-stage preclinical screening. The complete system is contained within a housing unit of approximately 1.9 m³.

Recent iterations of the system have also introduced substantial hardware and software advancements (Cliffe et al., 2023). These include the transition from a single-pump configuration to a centralised pump control module capable of regulating flow across 16 parallel units, enabling increased experimental scalability. The incorporation of automated live-cell imaging further enhances real-time monitoring capabilities, while improvements in system ergonomics (housing unit and loading of fluidic module) and graphical user interface design facilitate usability and experimental reproducibility.

Although significant progress has been made, further hardware and manufacturing refinements are ongoing. The Bioplate is currently fabricated from polymethyl methacrylate (PMMA), but future iterations will utilise injection-moulded polymeric materials such as polystyrene (PS) to reduce production time and improve optical clarity for imaging (Campbell et al., 2020). Planned redesign of well spacing will ensure compatibility with ANSI/SLAS 4-2004 standards for plate-handling technologies, while maintaining the existing ANSI/SLAS 1-2004 footprint. Bioplates are intended to be supplied as sterile, single-use consumables.

Additional developments include integration of a permanent injection and sampling port, enabling functions such as supernatant sampling and delivery of suspended cells, antibodies, growth factors, and other reagents during experiments. Ongoing optimisation of the fluidic module will also focus on material reduction to decrease weight and improve handling, particularly for routine transfer between incubators and laminar flow hoods. Future software development will incorporate automated beat counting, replacing the manual analysis used in this study and enabling more scalable data acquisition in line with the platform’s high-throughput design.

Taken together, these features position Mera as a modular and scalable platform that complements, rather than replaces, high-content cardiac MPS systems. It is particularly well suited to early-stage compound screening, combinatorial testing, and experimental designs requiring higher throughput, while more detailed functional cardiac characterisation remains better addressed by lower-throughput, sensor-integrated platforms.

## 5. Conclusion

Accurately predicting arrhythmogenic risk is a critical component of the drug development process. Conventional cardiac model systems rely on simple 2D culture formats, which have inherent limitations in mimicking the complex structure and function of *in vivo* tissue, and consequently, in predicting drug responses accurately. Our advanced microphysiological system (MPS), Mera, offers a robust solution by enabling the evaluation of beat-modulating drugs on cardiomyocyte and cardiomyocyte/cardiac fibroblast spheroids. With its modular design, automation features, and scalability to 640 spheroids, Mera offers a powerful and cost-efficient platform for preclinical cardiotoxicity screening, mechanistic drug evaluation, and personalised medicine applications.

While the current platform primarily supports beat-rate assessment rather than comprehensive electrophysiological characterisation, future iterations incorporating embedded sensing technologies, multi-organ integration, and increased cellular complexity are expected to further enhance its physiological relevance and predictive capability. This work makes a distinct contribution to the field of preclinical cardiac safety and efficacy testing by providing a scalable, semi-automated and physiologically relevant MPS capable of modelling drug induced modulation of beat rate in human-derived 3D cardiac tissues. By delivering human-relevant, reproducible, and high-throughput data while reducing reliance on traditional animal and low-complexity in vitro models, Mera represents a strong new approach methodology for predictive cardiac safety assessment. By bridging key translational gaps between traditional in vitro assays and in vivo outcomes, Mera offers a promising tool for predictive drug development.

## 6. Author Contributions

**Nuno Almeida:** Conceptualization, Writing - original draft, Methodology, Investigation, Formal analysis, Validation**; Finola Cliffe:** Conceptualization, Writing - original draft, Funding acquisition, Supervision, Project administration; **Mark Lyons:** Conceptualization, Funding acquisition, Supervision; **Veasna SumCoffey:** Writing – review & editing, Resources, Methodology, Investigation; **Sandra Sunil:** Writing - original draft, Methodology, Investigation; **Bhairavi Bengaluru Keshava:** Investigation; **Patrick Costello:** Methodology, Investigation, Visualisation, **Conor Madden:** Methodology, Investigation; **Shane Devitt:** Methodology, Investigation; **Sumir Ramesh Mukkunda:** Methodology, Investigation, Data curation, Software; **Seonaid Deely:** Methodology, Investigation, Visualisation; **Chiara Alessia de Benedictis:** Methodology, Investigation; **Leon G. Riley:** Methodology.

## 7. Data Availability Statement

The datasets generated and analyzed during the current study are not publicly available due to intellectual property and commercial confidentiality restrictions.

## 8. Statements and Declarations

All authors are employees or past employees of Hooke Bio, the developer of the Mera system described in this study. Some authors hold shares or stock options in the company. The authors declare no other competing interests.

